# Rapid inactivation of SARS-CoV-2 with ozone water

**DOI:** 10.1101/2020.11.01.361766

**Authors:** Hiroko Inagaki, Akatsuki Saito, Putu Eka Sudaryatma, Hironobu Sugiyama, Tamaki Okabayashi, Shouichi Fujimoto

**Author notes:** **Corresponding author:** Shouichi Fujimoto, Department of Hemovascular Medicine and Artificial Organs, Faculty of Medicine, University of Miyazaki, 5200 Kihara, Kiyotake, Miyazaki 889-1692, Japan.

## Abstract

Although ozone water is one of the promising candidates for hand hygiene to prevent fomite infection, the detailed effects of ozone water on SARS-CoV-2 have not been clarified. We evaluated the inactivating effect of ozone water against SARS-CoV-2 by its concentration and exposure time. The reduction rates of virus titer after 5 sec treatment with ozone concentrations of 1, 4, 7, and 10 mg/L were 81.4%, 93.2%, 96.6%, and 96.6%, respectively. No further decrease in virus titer was observed by the extended exposure time over 5 sec. High-concentration ozone water was considered to be effective in promptly inactivating SARS-CoV-2 virus.

## Introduction

Half a year has passed since the outbreak of the novel coronavirus disease 2019 (COVID-19) was first confirmed [1, 2], but COVID-19 continues to spread throughout the world, and the total number of patients approached 25 million as of August 2020 [3]. In addition to direct person-to-person transmission via respiratory droplets, exposure to contaminated material is considered to be an important transmission route [4, 5], because SARS-CoV-2 remains infectious for hours to days on contaminated surfaces or the human skin [6–9]. To prevent fomite infection, hand hygiene using various chemical compounds is recommended, especially in hospital and lavatory settings [9–11]. The United States Centers for Disease Control and Prevention (CDC) recommends hand hygiene with soap and water or alcohol-based hand rub (at least 60% alcohol), and that hand washing should be done for at least 40-60 seconds based on World Health Organization (WHO) recommendations [12]. A variety of disinfectants can reduce viral viability with a contact time of at least 30 seconds or more [10]. Among hand rub solutions, ozone water is one of the promising candidates, because it is known that ozone water has an inactivating effect on some microorganisms [13–16] and is harmless to human beings [17, 18]. Furthermore, the production of high-concentration ozone water has been facilitated by the advent of electrolytic ozone generators [19]. However, the details of the effects of ozone water on SARS-CoV-2 have not yet been clarified. In this study, the inactivating effect of ozone water on SARS-CoV-2 by concentration and contact time was evaluated.

## Materials and Methods

A strain of SARS-CoV-2 isolated from a patient who developed COVID-19 on the cruise ship *Diamond Princess* in Japan in February 2020 [20] was obtained from the Kanagawa Prefectural Institute of Public Health (SARS-CoV-2/Hu/DP/Kng/19-027, LC528233). The virus was propagated as described previously [21]. Ozone water was created by an electrolytic ozone water-generating device (Handlex, ONR-1, Nikkiso/Nikkamicron Co., Saitama/Tokyo, Japan) using sterile tap water. The concentration of ozone was measured by a portable analyzer (O3METER OZ-20, DKK-TOA CO., Tokyo, Japan).

To evaluate the antiviral effect of ozone water on SARS-CoV-2, aliquots of virus stock (10 μL) were put in the micro spitz. Ozone water (990 μL each) with a concentration of 1, 4, 7, or 10 mg/L was prepared and mixed with the above virus stock solution. Based on previous reports [14, 22], 100 μL of sodium thiosulfate solution (Na_2_S_2_O_3_) was added to each mixture of virus stock and ozone water at 5,10, or 20 sec (n=3) to terminate the ozone water reaction.

After adding Na_2_S_2_O_3_, the virus solutions were serially diluted in 10-fold steps using serum-free minimum essential medium (MEM) and then inoculated onto Vero cell monolayers in a 12-well plate. After adsorption of virus for 2 h, cells were overlaid with MEM containing 1% carboxymethyl cellulose and 2% FBS (final concentration). Cells were incubated for 72 h in a CO2 incubator, and then cytopathic effects were observed under a microscope.

Virus suspension mixed with sterile tap water without ozone was used as a negative control. To calculate plaque forming units (PFUs), cells were fixed with 10% formalin for 30 min, followed by staining with 0.1% methylene blue solution. The antiviral effects of ozone water were assessed using the logPFU ratio, as described previously [21]. All experiments were performed in a BSL-3 laboratory.

## Results

Ozone water at concentrations of 1, 4, 7, and 10 mg/L was prepared. Actual ozone concentrations were 1.0, 4.3, 7.4, and 10.3 mg/L, respectively. Na_2_S_2_O_3_ was used to stop the reaction. In SARS-CoV-2-infected cells, there was a marked cytopathic effect in cells inoculated with viruses without ozone water treatment (Fig. 1a). In contrast, cells inoculated with viruses treated with 7 mg/L ozone (Fig. 1b) showed largely comparable morphology to mock cells (Fig. 1c).

**Figure 1.**
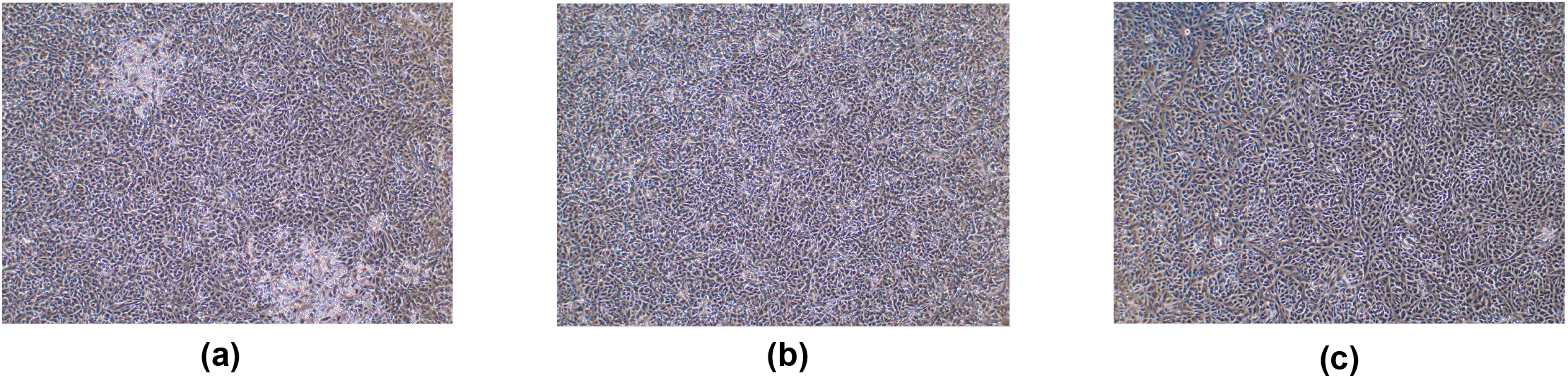
Cytopathic changes in SARS-CoV-2-infected Vero cells. (a) Tap water (0 mg/L, 20 sec), (b) ozone water (7 mg/L, 5 sec), and (c) mock-infected cells.

The dose- and time-dependent inactivating effect of ozone water on SARS-CoV-2 was evaluated. The plaque assay (Fig. 2) showed that ozone water inactivated SARS-CoV-2, but sterile tap water did not (Fig. 3). Specifically, the reduction rates of virus titer after 5 or 10-sec treatment for the concentrations (1, 4, 7, and 10 mg/L) of ozone water were respectively 81.4%, 93.2%, 96.6%, and 96.6% for 5 sec, whereas it was 75.4%, 93.2%, 96.6%, and 97.5% for 10 sec. There was a significant difference in the effect depending on the ozone concentration (Fig. 4). There was no significant difference between the effects of ozone water at concentrations of 7 mg/L and 10 mg/L. However, no further decrease in virus titer was observed even if the exposure time was extended to longer than 10 sec for any concentration of ozone water in the static state (Figs. 3). Taken together, ozone water showed a significant antiviral effect on SARS-CoV-2 in a dose-dependent manner.

**Figure 2.**
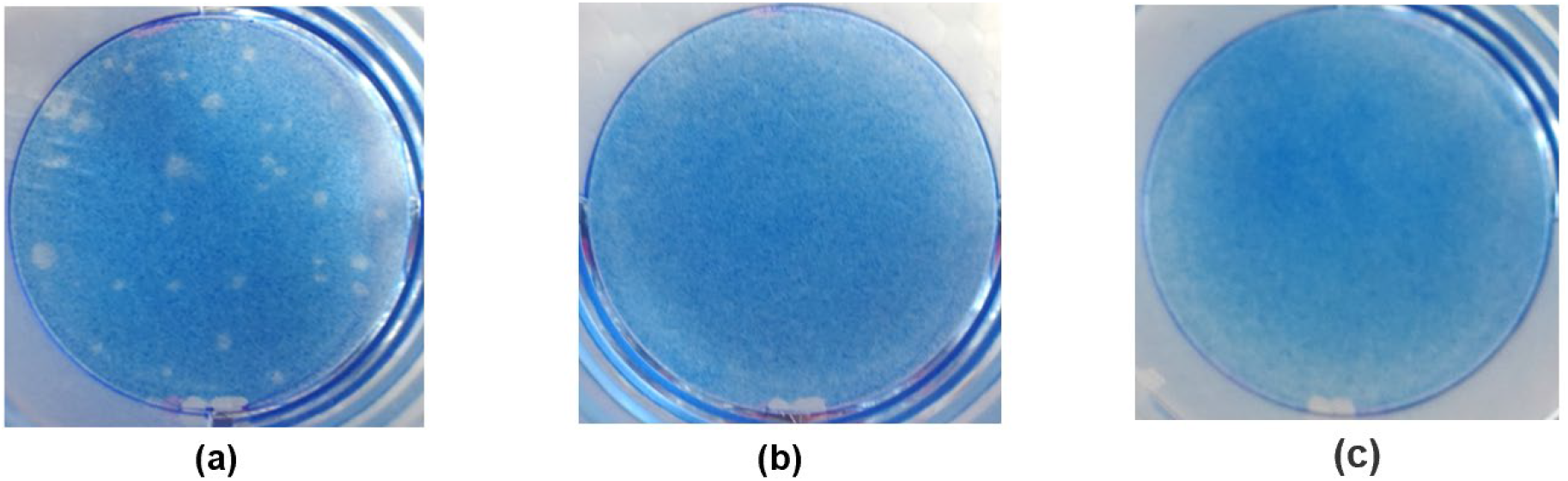
Plaque formation in Vero cells. Virus solutions were exposed to ozone water for 5 sec and subjected to a plaque assay. (a) virus and tap water (no ozone), (b) virus and 7 mg/L ozone water, and (c) mock-infected cells.

**Figure 3.**
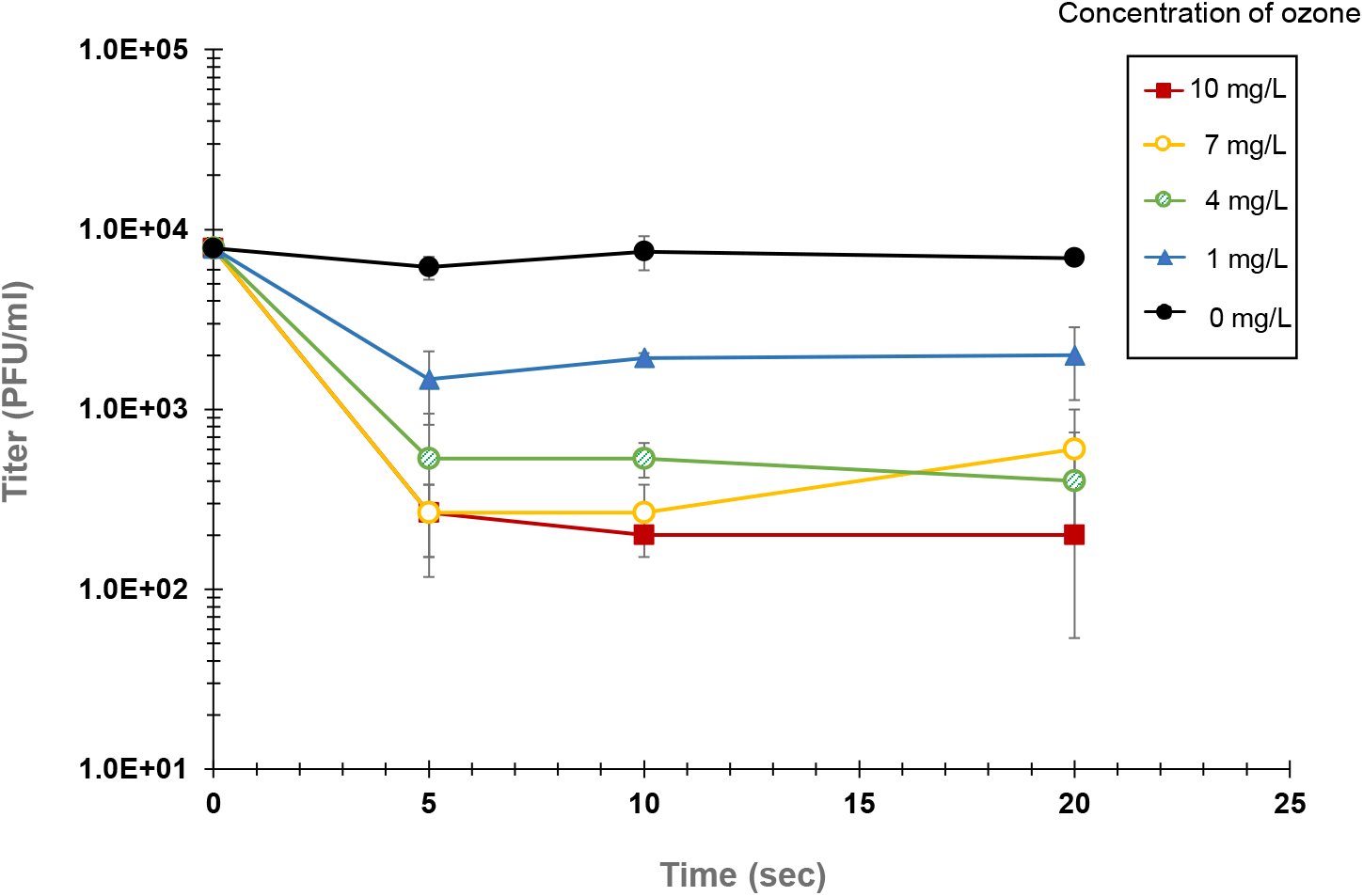
Longitudinal change of virus titers after treatment with ozone water.

**Figure 4.**
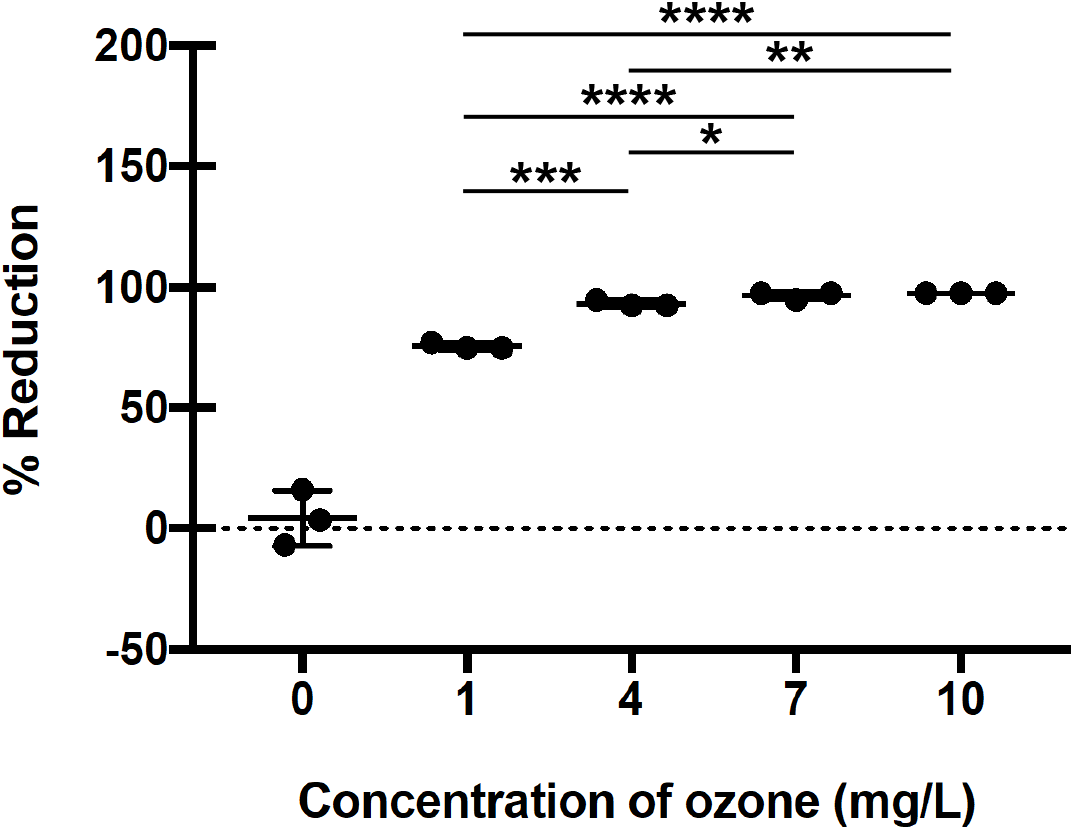
Reduction of virus titers by treatment with different concentrations of ozone water. Virus solutions were mixed with ozone water for 10 sec and subjected to a plaque assay. ****p < 0.0001, ***p < 0.001, **p < 0.01, *p < 0.05.

## Discussion

The aim of this study was to clarify the inactivating effect of ozone water on SARS-CoV-2 by its concentration and contact time. Although ozone water is known to have an inactivating effect on several microorganisms [13, 15, 16], whether ozone water is also effective against SARS-CoV-2 has not been reported so far. In addition, the mode of action of ozone water, including concentration and duration of treatment, should be investigated for its use to prevent fomite infection. This study showed that short time (5 sec) treatment with ozone water was sufficient to inactivate SARS-CoV-2. Recently, it has become possible to generate high-concentration ozone water due to the advent of electrolytic ozone generators. The present study demonstrated that high-concentration ozone water (7 mg/L, 10 mg/L) was more effective against SARS-CoV-2 than low-to medium-concentration ozone water (1 mg/L, 4 mg/L). Ozone water may show a limited effect on microorganisms depending on the conditions, but high-concentration ozone water (≥10 mg/L) has enough of an effect even in the presence of inhibitory materials such as proteins [16]. Although medium-concentration (4 mg/L) ozone water may be equivalent or slightly inferior to propanol (60%)-based hand rubs for disinfecting hands [23], one of the advantages of ozone water is that it causes marginal or no skin damage when disinfecting hands [17, 18]. Considering these issues, the fact that higher concentrations of ozone water show stronger inactivating effects on SARS-CoV-2 may be a useful finding when using ozone-based hand disinfectants to prevent fomite infection.

Interestingly, there was no time-dependent inactivating effect of ozone water on SARS-CoV-2. One possible explanation for this phenomenon is that the inactivating effect was already achieved after 5-10 sec of exposure with all concentrations of ozone water. Alternatively, the detection limit of the plaque assay might lead to this result, at least for 7 or 10 mg/L ozone water. Another explanation is that the ozone activity was quickly attenuated by culture medium containing 2% FBS in all concentrations of ozone water. To test this possibility, the decrease of ozone concentration after adding the culture medium was examined (Supplementary Fig. 1). There was marked attenuation of the ozone concentration, as it was halved in about 10 seconds in all grades, especially at the low ozone concentration (1 mg/L). A few reports have shown the time-dependent antimicrobial activity of ozone water [22, 24]. However, there are some confounding factors (type or strain of microorganism, methods of measuring antimicrobial efficacy and ozone concentration, presence of ozone-demanding medium components, and so on) that make it difficult to compare the results of each study.

Despite the significant inactivating effect of ozone water, the present study may have some limitations. First, ozone concentrations measured in this study may not reflect the exact value of ozone water in the microtube contacting the virus, since it is known that the ozone concentration decreases rapidly in the environment. Given that this study focused only on SARS-CoV-2, the inactivating effect of ozone water on other microorganisms needs to be investigated in future studies. Furthermore, since the results of this experiment were obtained in a static state (not dynamic) rather than in running water, different outcomes may be observed when these two conditions are compared. In future studies, more practical information can be obtained by evaluating the ozone water concentration and exposure time for the optimal inactivating effect under various conditions.

Currently, we are faced with the need to prevent SARS-CoV-2 infection for a long time, and a new lifestyle, so called “with corona”, is needed worldwide. Washing hands is a critical way to prevent fomite infection, as suggested by the WHO. Very recently, the virucidal efficacy of formulated microbicidals, such as ethyl alcohol, para-chloro-meta-xylenol, and quaternary ammonium compounds, has been reported for inactivating SARS-CoV-2 [25]. On the other hand, it has been pointed out that SARS-CoV-2 may survive for 9 hours on the skin [9]. In addition, most of the touchable surfaces in a designated hospital for COVID-19 were found to be heavily contaminated, suggesting that environmental surfaces are a potential medium of disease transmission [26]. Hygiene of exposed skin with high-concentration ozone water, which is less likely cause skin damage, will be a useful and sustainable way for person with sensitive skin to contain the spread of SARS-CoV-2 infection.

## Acknowledgements

The authors would like to thank Drs. Tomohiko Takasaki and Jun-Ichi Sakuragi, from the Kanagawa Prefectural Institute of Public Health, for providing the SARS-CoV-2/Hu/DP/Kng/19-027 strain.

## Conflict of interest statement

H.S. receives part of his salary from Nikkiso Co., Ltd., Tokyo, Japan. Nikkiso had no role in study design, data collection and analysis, decision to publish, or preparation of the manuscript. The other authors declare no conflicts of interest.

## Funding sources

None.

## Contributors

H.I. and H.S. conceived the study. H.I. and A.S. wrote the manuscript. A.S., S.PK, and T.O. conducted the experiments dealing with viruses. S.F. contributed to the study design, study supervision, and manuscript revision.

---

**Supplementary Figure 1.**
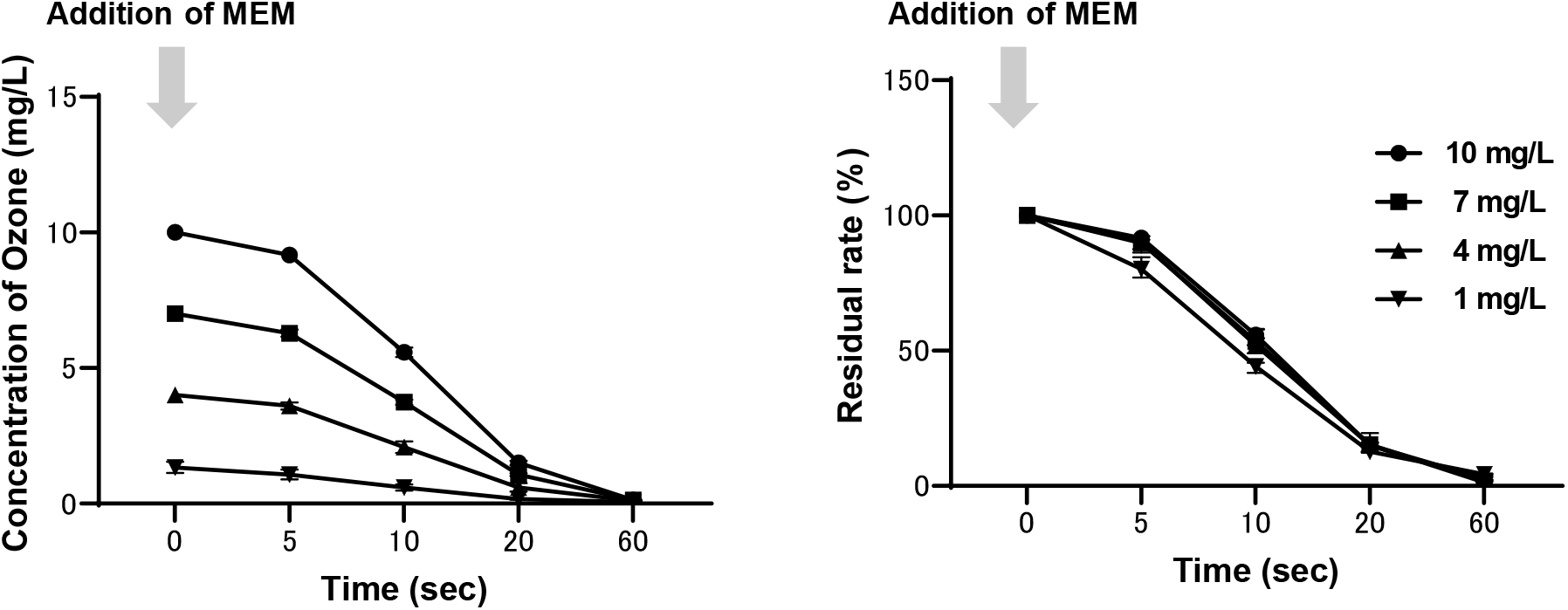
Rapid decrease of ozone concentration by the addition of culture medium (MEM, 1 mL) for each concentration of ozone water (100 mL). Lt; Concentration of ozone (mg/L), Rt; Residual rate (%).

